# A Chromatin Biology Assessment of AlphaFold3

**DOI:** 10.64898/2026.06.26.734680

**Authors:** Yash Bhargava, Cynthia Wolberger, Sanim Rahman

## Abstract

Biomolecular structure prediction tools such as AlphaFold have achieved remarkable success in predicting structures of single proteins and multiprotein complexes. AlphaFold3 now incorporates the capability to model complexes containing nucleic acids and chemically modified side chains. Investigators can now predict structures of proteins bound to chromatin, where interactions with nucleosomal DNA and histone post-translational modifications converge to control genome function. To evaluate its robustness in modeling chromatin complexes, we benchmarked AlphaFold3 on 115 structures containing nucleosomes whose coordinates were released by the Protein Data Bank after the training set cutoff date. We find that AlphaFold3 excels at predicting histone-driven interactions and accurately models complexes that deposit and recognize post-translational modifications. By contrast, AlphaFold3 struggles to predict structures of chromatin factors that primarily engage nucleosomal DNA, notably transcription factors and chromatin remodelers. Finally, we show that AlphaFold3 can faithfully recapitulate known post-translational modification recognition patterns, matching experimentally determined specificity profiles. This assessment of the capabilities and limitations of AF3 in chromatin structural biology provides a roadmap for its effective application to studies of chromatin regulation and PTM readout, while identifying key areas for future algorithmic refinement.

**Significance:** Structure prediction with AlphaFold has become an invaluable tool in experimental biology, and the accuracy of many of its predictions has been verified in structural and biochemical studies. With the recent incorporation into AlphaFold3 of nucleic acids and post-translational modifications, this prediction tool can now be applied to chromatin structural biology. Our benchmarking of AlphaFold3 reveals its strengths and weaknesses in predicting structures of proteins bound to nucleosomes, thereby providing a framework for using these models in mechanistic studies of chromatin regulation. We introduce metrics for evaluating structures of nucleosome complexes that highlight AlphaFold3’s strengths in predicting protein-nucleosome interactions and post-translational modification specificity.

## Introduction

Eukaryotic genomes are packaged into chromatin, a nucleoprotein complex whose fundamental organizational unit is the nucleosome. Each nucleosome contains ∼147 bp of DNA wrapped around an octameric protein core containing two copies each of histones H2A, H2B, H3, and H4. Virtually all cellular processes requiring access to DNA – including transcription, DNA replication, and DNA repair – depend upon protein complexes that interact with nucleosomes.

Post-translational modifications (PTMs) of the core histone proteins, such as acetylation, methylation, phosphorylation and ubiquitination, play a central role in regulating these processes by facilitating or inhibiting recruitment of chromatin factors (1, 2), as well as by altering the higher-order packaging of the genome (3–5).

Our understanding of how protein complexes engage nucleosomes has greatly increased since the first high-resolution structure of a nucleosome was reported in 1997 (6). The determination of a handful of crystal structures of proteins bound to nucleosomes (7–11) over the next 15 years provided the first insights into the mechanism by which proteins were recruited to chromatin.

The 2014 “resolution revolution” in cryogenic electron microscopy (cryo-EM) dramatically increased the number and complexity of structures of chromatin complexes, which had been difficult to crystallize and were often available in very limited amounts. By 2025, there were 798 nucleosome-containing complexes deposited in the Protein Data Bank (PDB), largely determined by cryo-EM.

The advent of AlphaFold2 (12), a powerful deep learning tool for protein structure prediction, transformed structural biology by providing accurate models that have facilitated interpretation of low-resolution cryo-EM maps and generated testable mechanistic hypotheses (13–18).

However, the inability to incorporate DNA or post-translational modifications in the predictions made it impossible to model complexes with full nucleosomes or to predict the impact of PTMs on complex formation. The recent introduction of AlphaFold3 (AF3) (19), which integrates proteins, nucleic acids, small molecules, ions, and PTMs within a unified modeling framework, now makes it possible to model complexes containing nucleosomes. There are examples of recent structures for which AF3 successfully predicted features such as recognition of the nucleosome acidic patch (15, 20, 21). However, the accuracy of AF3 in modeling complexes that contain chromatin has not been systematically evaluated.

To assess the ability of AF3 to accurately model the architectural and regulatory complexity of protein complexes bound to nucleosomes, we systematically benchmarked AF3 against 115 structures containing nucleosomes that were deposited in the Protein Data Bank (PDB) after the training set cutoff date. By evaluating predicted models using interface-specific metrics, we identify contexts in which AF3 performs accurately and those where its predictions falter. We find that AF3 performs well in predicting the structure of complexes that interact primarily with the octameric histone core of the nucleosome, whereas its performance is significantly worse for protein complexes that interact primarily with the nucleosomal DNA. In addition, we find that AF3 can accurately recapitulate PTM readout, correctly predicting the specificity reported in experimental screens. Our findings provide insights into the current capabilities and limitations of deep learning-based structure prediction in chromatin biology and establish a foundation for its effective application in studying chromatin regulation and PTM readout.

## Results

### Overall model confidence reveals AF3’s accuracy in mono-nucleosome structure and weaknesses in multi-nucleosome complexes

To evaluate the ability of AF3 to accurately predict structures of complexes containing nucleosomes, we curated a benchmark dataset (Figure 1) of complexes released after the AF3 training cutoff (September 30^th^, 2021) and until September 1^st^, 2024 (19). We removed structures released after the cutoff that had already been deposited in the PDB prior to the cutoff date, complexes over the AF3 server limit of 5,000 tokens, and complexes containing DNA damage sites, which cannot be modeled in AF3 (see Methods). These criteria refined our initial list of 322 structures released after the training cutoff to 115 structures, which we used for systematic benchmarking. Given the broad range of functions and nucleosome-binding mechanisms represented in our benchmarking set, we classified the 115 structures into distinct groups that would reflect the ability of AF3 to predict protein-protein, protein-DNA, and protein-PTM interactions. These classes were then further categorized according to their biological function and/or enzymatic activity. Predictions of structures containing nucleosomes alone were grouped as mononucleosomes (n=23), multi-nucleosomes (n=13), and subnucleosomes (n=6), the latter containing less than the full histone octamer. The remaining complexes were classified as PTM writers (enzymes that modify histone residues; n=21), PTM readers/erasers (proteins that bind to PTMs and either recognize or remove the modification; n=13), histone variant readers (n=5), chromatin remodelers (n=11) and transcription factors (n=18). Structures that could not be grouped in these categories are binned as other nucleosome interactors (n=15).

**Figure 1.**
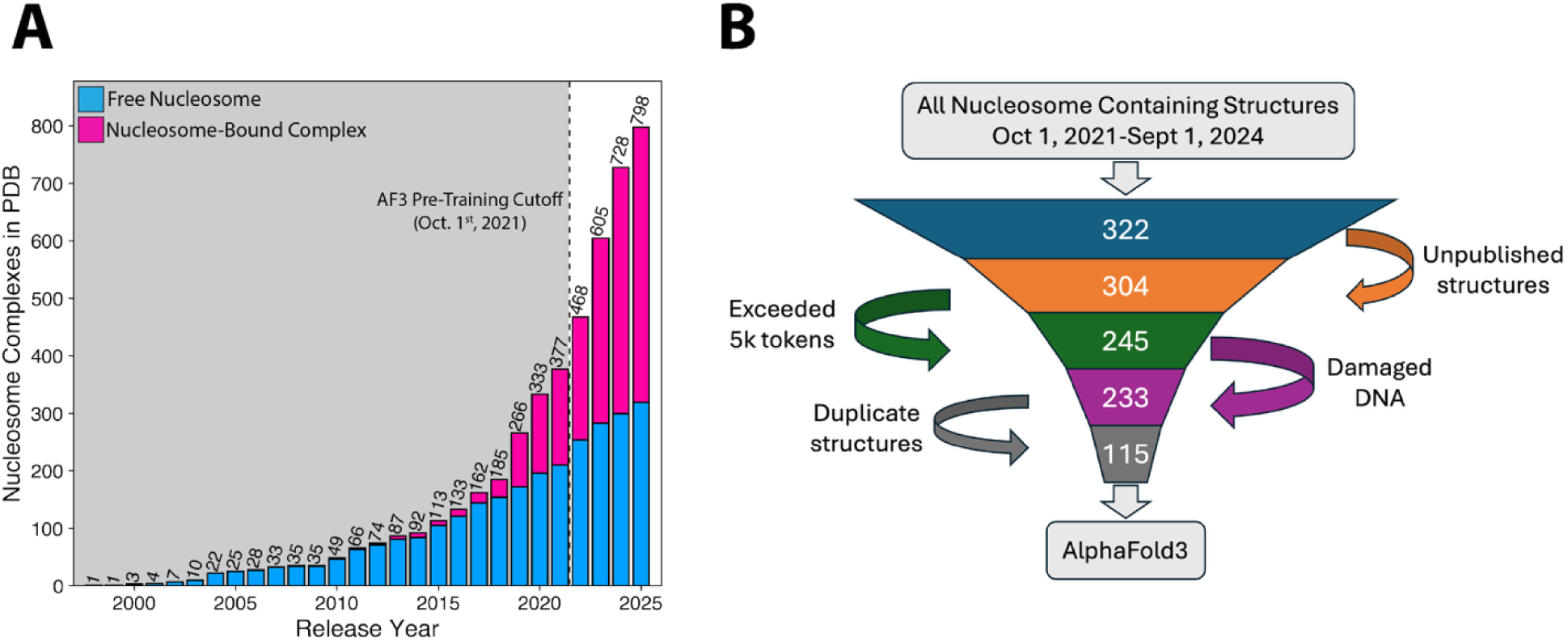
Curation of nucleosome benchmark dataset. (A) Number of nucleosome and nucleosome-like structures deposited in the Protein Data Bank (PDB) each year solved with or without a nucleosome-binding protein from NucleosomeDB (54). The vertical dashed line depicts the AlphaFold3 (AF3) training dataset cutoff date. (B) Dataset curation of all structures released after the AF3 training cutoff for evaluating nucleosome and nucleosome-interactor structure prediction.

All AF3 predictions were performed using the publicly available server with the default settings. For entities that that are not supported on the AF3 server, including some ligands, ions and selected chemical modifications of proteins and DNA, ligands and ions we made substitutions or deletions summarized in SI Table 1 (further prediction details described in Methods). However, in the case of ubiquitin modifications, which are not supported by the AF3 server, we used AlphaFold3X (22) to incorporate an explicit disulfide crosslink between the C-terminal residue of ubiquitin and modified histone residues. This approach to modeling ubiquitinated lysines was recently reported to improve the prediction of ubiquitinated proteins and poly-ubiquitin (23) (see Methods).

We first evaluated the confidence of the entire predicted assembly using the interface predicted Template Modeling (ipTM) score, which reflects the confidence of the predicted interaction interfaces within the model (Figure 2A). Out of the 115 models, 27 models had an ipTM of ≥0.80, indicating a high-confidence model, while 37 models had an ipTM of 0.60-0.79, which has been referred to as a “grey area” in which predictions could be either correct or wrong (24). The remaining 51 models had ipTM scores below 0.60, which are considered failed predictions (24).

**Figure 2.**
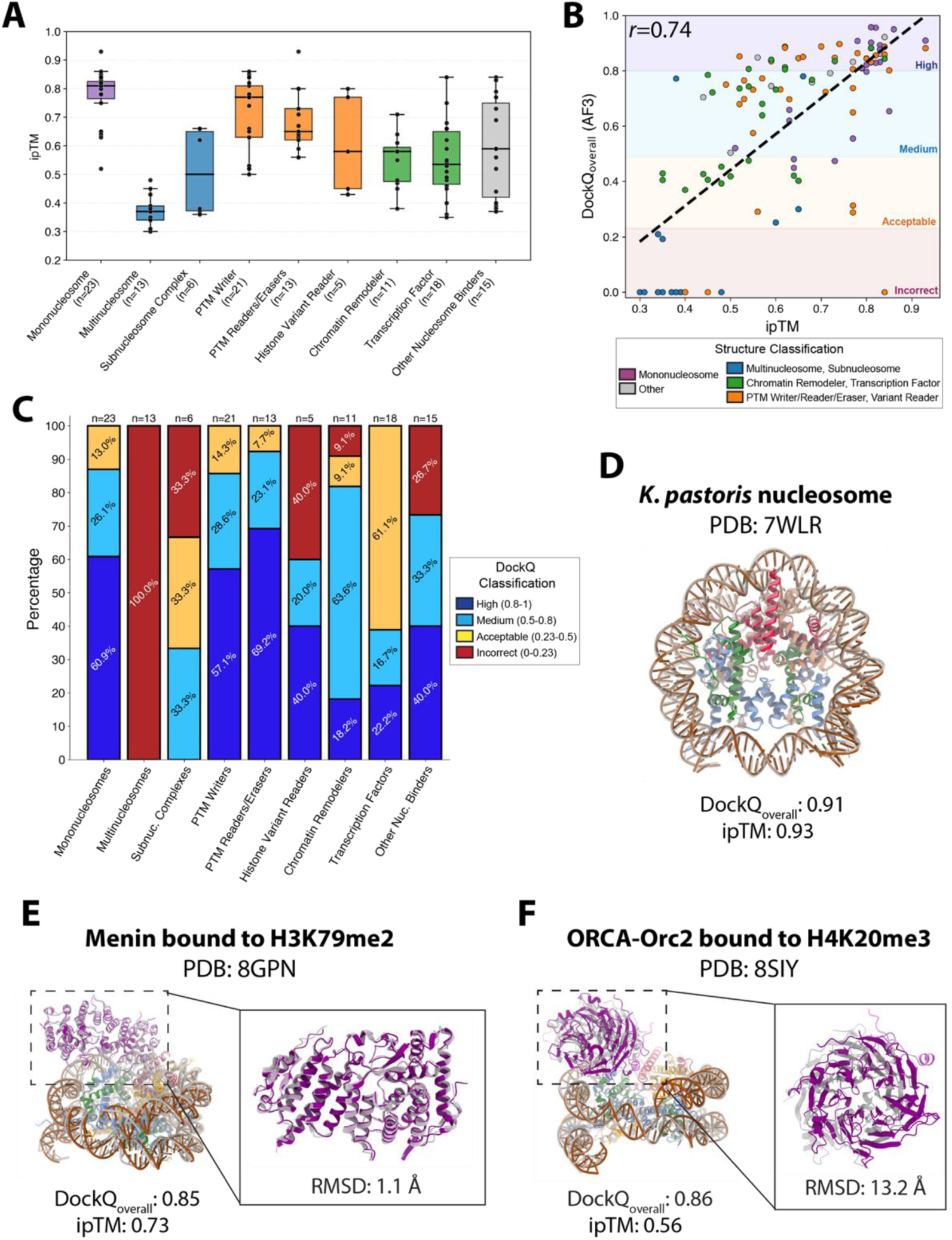
Performance of AlphaFold3 (AF3) on predicting overall nucleosome and nucleosome-interactor structures. (A) Distribution of interface predicted Template Modeling (ipTM) score for all benchmarked nucleosome- and nucleosome-interactor structures classified by their biological function, enzymatic activity, or assembly. (B) Correlation between DockQ_overall_ score and ipTM score for all benchmarked structures. (C) DockQ_overall_ distribution for all benchmarked structures classified by their biological function, enzymatic activity, or assembly. (D) Cryo-electron microscopy (Cryo-EM) structure of the *K. pastoris* nucleosome (gray) overlaid on the AF3 predicted model (colored). Cryo-EM structure comparison of the experimental (gray) and AF3 (colored) model of (E) Menin bound to a H3K79me2 nucleosome and (F) ORCA-Orc2 bound to H4K20me3 nucleosome with the RMSD of Menin and ORCA-Orc2 calculated based on the alignment of the nucleosome.

Although the ipTM score provides a valuable assessment of biomolecular interaction confidence, there are two significant drawbacks to interpreting this score as reflecting the accuracy of biomolecular interactions. First, the presence of disordered regions and/or accessory domains that do not contribute to protein binding negatively impact ipTM (25, 26). This is a particular issue for structures of proteins bound to nucleosomes, as the presence of flexible histone tails decreases ipTM despite not being involved in the direct interaction (SI Figure 1). Second, ipTM only reflects the confidence of AF3 in the overall prediction, not how well the prediction compares to a reference structure.

We used DockQ_overall_ (27) to compare all predicted complexes to the experimentally determined PDB structure. The DockQ_overall_ score combines several measures of model quality and has proved more robust in evaluating structure predictions (27–29). In short, DockQ_overall_ reports the accuracy of a predicted model against an experimental reference structure by assessing the similarity in all protein-protein, protein-nucleic acid, and nucleic acid-nucleic acid interaction interfaces. For all multi-nucleosome models and some poorly predicted complexes (n=17), DockQ_overall_ analysis was not possible due to severe prediction inaccuracies that prevented initial structural alignment to the reference experimental structure, consistent with the observed poor ipTM scores (Figure 2A). These models were therefore not further analyzed or considered. To ensure that we could evaluate the highest quality structure prediction, we computed DockQ_overall_ scores for all five models output by AF3 for each structure prediction and used the model with the highest DockQ_overall_ for downstream analysis. Among the models that could be evaluated, the overall DockQ_overall_ score is strongly correlated with ipTM (*r*=0.74) (Figure 2B). Based on the DockQ_overall_ score, AF3 was able to predict 45/115 models within the “High” accuracy (0.8-1.0) range and 31/115 were within “Medium” accuracy (0.5≤DockQ<0.8). On the lower end of prediction accuracy, 20/115 scored within the “Acceptable” accuracy range (0.23≤DockQ<0.5) and 19/115 in the “Incorrect” accuracy range (DockQ<0.23 or failed DockQ calculation).

The great majority of structure predictions that had both an excellent ipTM score (ipTM ≥0.8) and a DockQ_overall_ score in the “High” accuracy distribution were of isolated mono-nucleosomes, including histone variants, histones from various organisms, or non-Widom 601 DNA sequences (Figure 2C). This is perhaps not surprising, since the histone octamer core of the nucleosome is highly conserved in structure and there are hundreds of nucleosome structures in the training data set. By contrast, AF3 had low confidence metrics for predictions of non-canonical nucleosome structures such as tetrasomes and hexasomes, or multi-nucleosome structures (Figures 2A, 2C), as the majority of models had an ipTM score below 0.6 and/or DockQ_overall_ scores below 0.23 or were uncalculatable due to severe prediction inaccuracy.

A comparison of predicted versus experimentally determined complex structures revealed that the ipTM and DockQ_overall_ score were an inconsistent measure of prediction accuracy in many cases. Nearly half of the models were predicted with an ipTM≥0.6 and a DockQ_overall_ score that spans the Medium and High confidence classes (n=53). Of these, the majority were interactor-bound nucleosome complexes. Upon visual inspection, we found that some predictions matched the experimentally determined structures well, whereas others had significant deviations in the orientation of the interactor on the nucleosome surface (Figure 2E-F). For example, predicted models of both the Menin-H3K79me2 (30) and ORCA-Orc2-H4K20me3 (31) nucleosome complexes were assigned a DockQ_overall_≥0.8, indicating high model accuracy. However, whereas the orientation of Menin on the nucleosome matches that in the reported cryo-EM structure (Figure 2E), with a RMSD of 1.1 Å, the orientation of ORCA-Orc2 on the nucleosome deviates substantially from the cryo-EM structure, with an RMSD of 13.2 Å from the cryo-EM structure (Figure 2F).

### Interactor-Nucleosome DockQ score highlights AF3’s strength in histone-driven interactions

We speculated that the discrepancy between DockQ_overall_ and the orientation of interactors on the nucleosome was due to contributions to the overall score by the histones in the octameric core of the nucleosome. DockQ_overall_ is calculated using a pairwise fraction of native contacts (f_nat_), scaled interface RMSD (iRMSD), and scaled ligand RMSD (LRMSD) (Equation 1), where the ligand is defined as the smaller chain (defined by number of amino acids or nucleotides) of each pairwise interaction (28, 29).

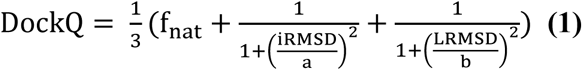

*f_nat_* is defined as the fraction of correctly predicted native contacts between separate protein or nucleic acid chains with at least one pair of heavy atoms that are within 5 Å of each other. iRMSD is the backbone (protein backbone is defined as N, CA, C, and O atoms; nucleic acid backbone is defined as P, OP[1,2], O[2-5], and C[1-5]) root-mean-square deviation of the interface residues between the predicted and reference complex, after superposition of the interface residues. LRMSD is the backbone root-mean-square deviation of the ligand with respect to the reference complex, after alignment to the receptor; *a* and *b* are the pre-defined scaling factors for iRMSD and LRMSD (1.5 and 8.5, respectively). In multi-chain predictions, these pairwise interactions (*M*) are amalgamated into an overall DockQ score which is defined as an unweighted average of all DockQ scores (Equation 2).

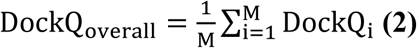

In the simple case of a single protein bound to a nucleosome, 55 pairwise interactions are considered, assuming all possible pairwise interactions satisfy the requirements for *f_nat_*. Out of the 55 interactions, 45 are within the nucleosome core particle alone. As a result, the DockQ_overall_ score largely reflects the high accuracy of the predicted nucleosome structure rather than of the pose of the interactor on the nucleosome.

To address the shortcoming of DockQ_overall_ in scoring predictions of how protein interactors interact with the nucleosome, we partitioned DockQ_overall_ scores to extract features specific to the interactor-nucleosome interaction (Figure 3A). For all interactor-containing models, we extract all pairwise DockQ calculations between the interactor and the histones or the DNA. These are defined as interactor-histone DockQ_Int,Hist_ (Equation 3) and interactor-DNA DockQ_Int,DNA_ (Equation 4).

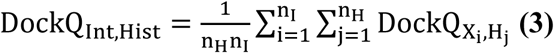

**Figure 3.**
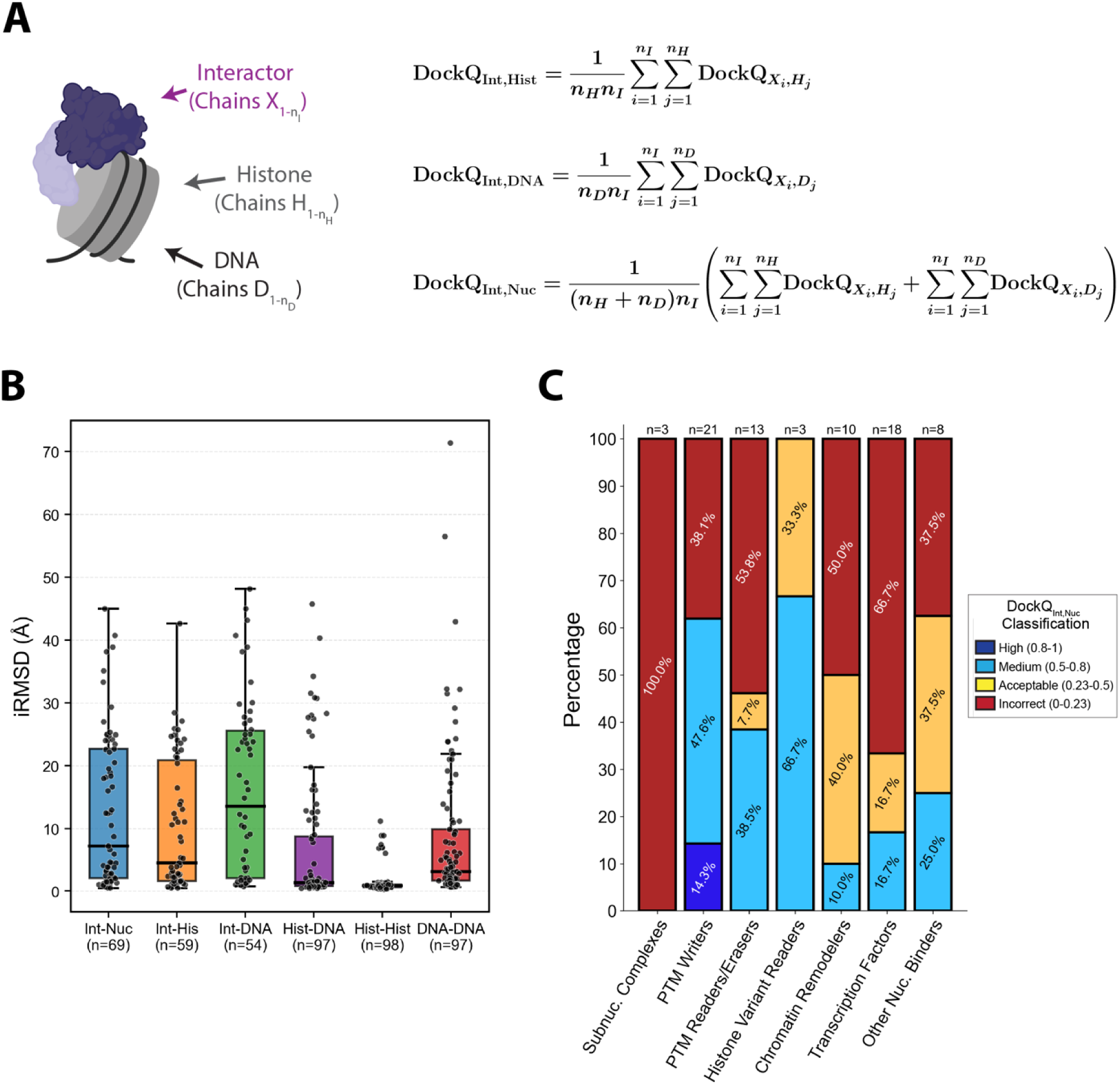
Calculation of DockQ_Int,Nuc_ score and analysis of nucleosome-interactor dataset. (A) Description and calculation of DockQ_Int,Hist_, DockQ_Int,DNA_, and DockQ_Int,Nuc_ scores. (B) Distribution of interface Root Mean Square Deviation (iRMSD), in angstroms, of all chain-chain interactions. (C) Distribution of DockQ_Int,Nuc_ scores based on the biological and enzymatic function of the nucleosome interactor.

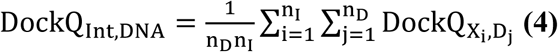

As a weighted average, the derivations can be used to yield DockQ_Int,Nuc_ (Equation 5).

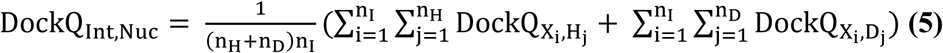

Since DockQ depends on iRMSD (Equation 1), we first assessed the model accuracy of different chain-chain interactions based on iRMSD (Figure 3B). As expected, we found that the pairwise interactions between histones in the nucleosome core had the lowest iRMSDs (median iRMSD=1.0 Å). On the other hand, we found that the overall iRMSD for interactor relative to nucleosome indicated quite variable prediction accuracy. However, when contacts of the interactor with histones and with DNA were considered separately, we found that contacts of interactors with the histones alone were predicted more accurately, with a median iRMSD of 4.6 Å, while pairwise interactor-DNA interactions were substantially worse, with a median iRMSD=13.6 Å. This discrepancy in interactor-histone and interactor-DNA accuracy is also reflected when assessing DockQ_Int,Nuc_ scores based on biological function of the interactor (Figure 3C). Interactors that primarily engage the histone octamer, such as histone writers, performed relatively well, with 61.9% (13/21) of predictions scoring within the “Medium” range or better. On the other hand, interactors that rely predominantly on DNA-driven interactions such as chromatin remodelers and transcription factors performed worse, with 10.0% (1/10) and 16.7% (3/18) of models scoring in the “Medium” range or better, respectively. These results suggest AF3 performs much better on complexes in which most nucleosome contacts are with the histone core.

### Predictions of interactions with nucleosomal DNA perform poorly in AF3

To better understand why AF3 did not perform well in predicting interactions with DNA, we examined interactors that predominantly contact nucleosomal DNA. Structures in this category are primarily transcription factors or chromatin remodelers. In the release of AF3 (19), the authors noted that a limitation of the model is that coordinates of protein-nucleic complexes that are both greater than 100 nucleotides and contain at least 2,000 residues in total contain clash violations. This limitation alone cannot account for the observed prediction inaccuracy, since 41 of the 58 predictions classified as Incorrect or Acceptable based on DockQ_Int,Nuc_ contained less than 100 nucleotides and 2,000 residues total.

Although some binders that form extensive contacts with nucleosomal DNA were predicted accurately, as reflected in high DockQ_Int,Nuc_ scores and agreement with experimental models (Figure 4A-B), we identified two classes of structure prediction errors. One class orients the transcription factor correctly on DNA but at the incorrect base sequence. MYC-MAX, a basic helix-loop-helix heterodimeric transcription factor, was positioned correctly on the nucleosomal DNA when the MYC-MAX binding motif was located at SHL +5.8 (PDB: 8OTT) (Figure 4A) (32). However, if the MYC-MAX recognition site was located at a position that is buried when the DNA is fully wrapped around the histone core, AF3 does not predict DNA unwrapping. Instead, the predicted structure positions MYC-MAX at SHL +5.1, a DNA site that is fully accessible but does not contain the cognate recognition sequence (Figure 4C). Notably, this structure (PDB: 8OTS) also contains a second bound transcription factor, OCT4, whose binding to an exposed cognate site at the other end of the DNA is correctly predicted (Figure 4C).

**Figure 4.**
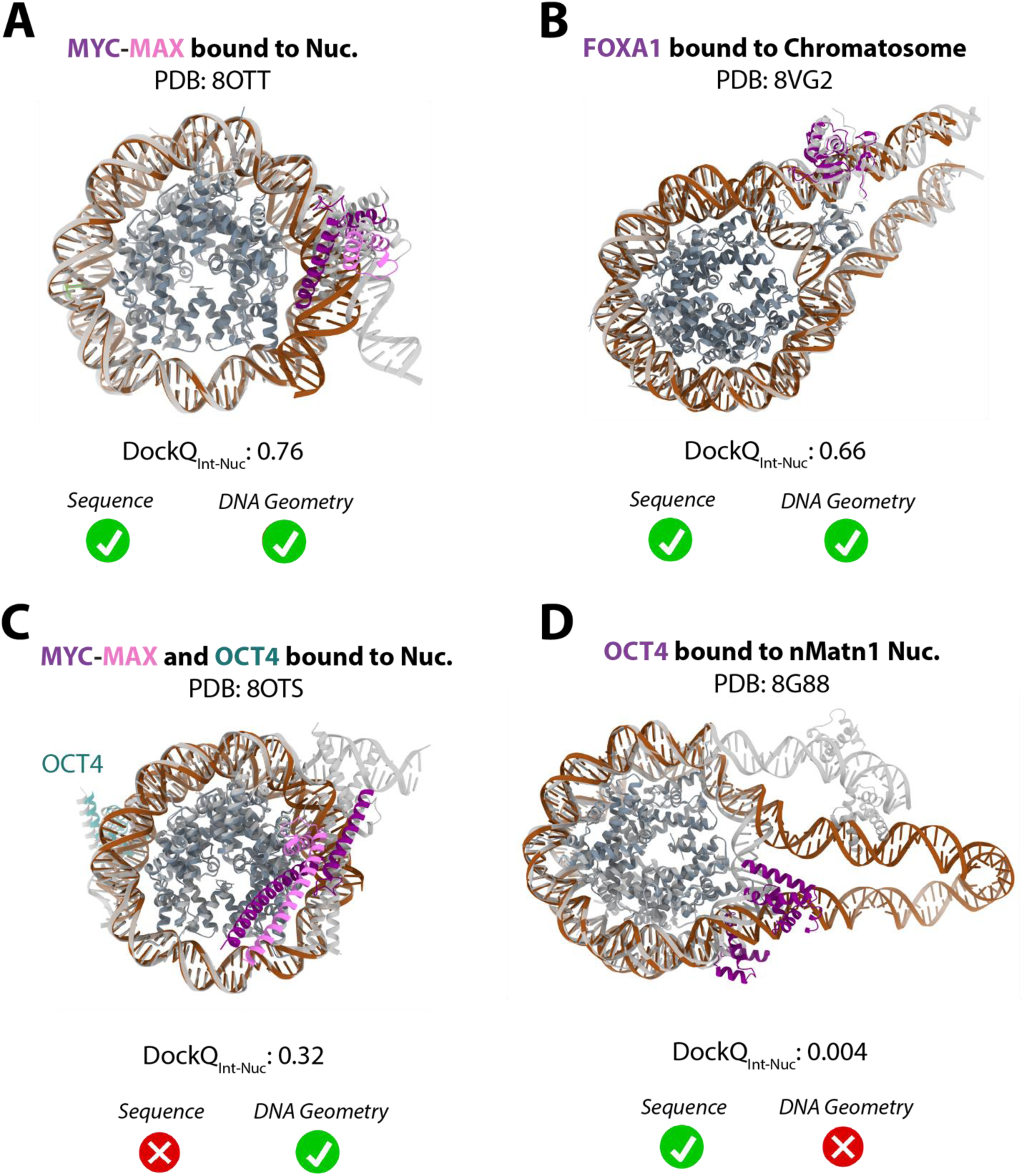
AlphaFold3 (AF3) predictions of transcription factors bound to nucleosomes. Examples of accurate transcription factor predictions of (A) MYC-MAX bound to a nucleosome (32) and (B) FoxA bound to a chromatosome (55). The AF3 predicted model (colored) is superimposed on the experimental structure (transparent gray). Examples of inaccurate transcription factor predictions of (C) MYC-MAX and OCT4 bound to a nucleosome (32) and (D) OCT4 bound to a nucleosome containing a nMatn1 DNA sequence (33).

Although the prediction results with MYC-MAX might suggest that AF3 always favors fully wrapped DNA, another class of transcription factor prediction errors provides striking counter-examples. In some cases, the AF3 prediction completely unwound a central stretch of the nucleosomal DNA to form a physically implausible loop that does not contact the histone octamer. An example is seen in the predicted structure of OCT4 bound to a nMatn1 nucleosome (PDB: 8G88) (Figure 4D) (33). Whereas the experimentally determined complex contains a DNA linker extension to which Oct4 binds, the AF3 prediction contains a long central loop of DNA that extends away from the histone core (Figure 4D). Interestingly, the DNA sequence to which OCT4 is predicted to bind is consistent with the experimental structure despite the severe errors in DNA structure (33). Together, our results suggest that AF3 suffers from limitations in predicting interactors that predominantly rely on nucleosomal DNA binding.

### AF3 can accurately predict the binding orientation of histone PTM writers and readers

Based on the DockQ_Int,Nuc_ distribution, AF3 performed particularly well on interactors that are histone PTM enzymes (“writers”), enzymes that remove these PTMs (“erasers”), and protein domains that recognized histone PTMs (“readers”) (Figure 3C). We therefore explored whether AF3 could model molecular interfaces that were critical for PTM readout or modification with residue-level accuracy. We selected models that scored well based on DockQ_Int,Nuc_ (Figure 3) and compared the prediction with available biochemical data. We first examined the interaction interface reported for Eaf3 bound to a nucleosome containing trimethylated histone H3-K36 (H3K36me3) (PDB: 7YI1) (Figure 5B) (34). The structure contains two symmetrically bound copies of Eaf3, each bound to one of the H3K36me3 histone tails. The AF3 prediction shows the same symmetric binding mode. In addition, the AF3 model recapitulates recognition of H3K36me3 by the Eaf3 aromatic cage through π-cation interactions (Figure 5B). Eaf3 also interacts with the nucleosomal DNA with residue K85, which is captured within the AF3 model (Figure 5B).

**Figure 5.**
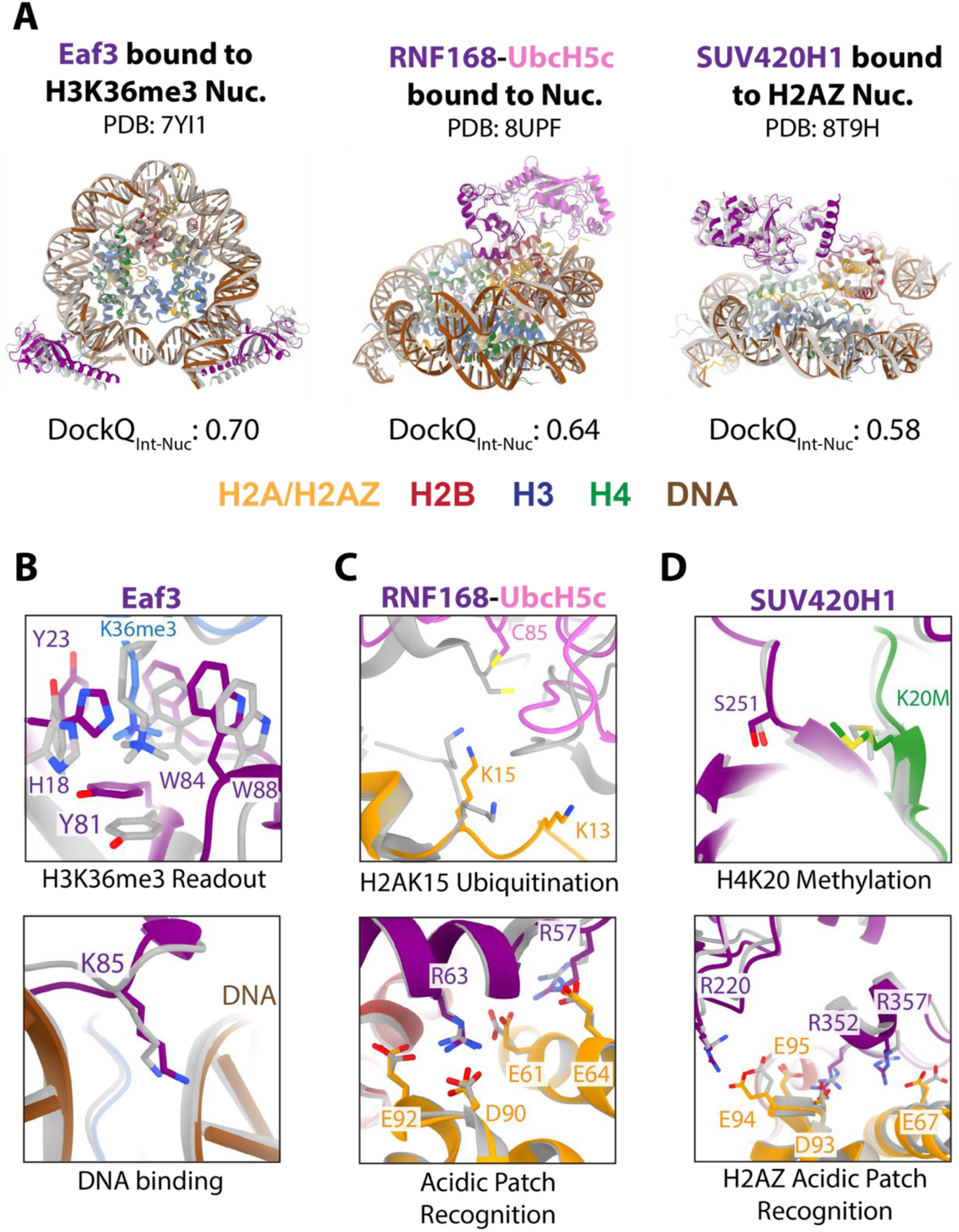
Accurate AlphaFold3 (AF3) predictions of post-translational modification (PTM) reader and writer binding to nucleosomes. (A) Overview of AF3 structure predictions of Eaf3 bound to a H3K36me3 nucleosome (34), RNF168-UbcH5c bound to an unmodified nucleosome (35), and SUV420H1 bound to a H2AZ nucleosome (36). The AF3 predicted model (colored) is superimposed onto the experimental structure (gray). (B) Zoom in on the recognition of H3K36me3 by the Eaf3 chromodomain and electrostatic interaction with the nucleosome DNA. (C) Close up view of the UbcH5c active site C85 residue in relation to histone H2A K13 and K15 and contacts between RNF168 and the nucleosome acidic patch. (D) Zoom in on the SUV420H1 active site S251 residue in relation to histone H4K20M and contacts between SUV420H1 with the H2AZ-containing nucleosome acidic patch.

An example of a histone writer whose predicted orientation on the nucleosome closely resembles the experimentally determined structure is UbcH5C-RNF168 (PDB: 8UPF) (35), an E2-E3 enzyme pair that ubiquitinates histone H2AK13 and H2AK15. In the AF3 model, both UbcH5C and RNF168 are positioned on the nucleosome surface in a manner similar to the reported cryo-EM structure (35), forming interactions with the nucleosome acidic patch (Figure 5A,C) and with the nucleosomal DNA (SI Figure 2A). The position of the UbcH5C catalytic cysteine (C85) relative to the target H2AK13 and H2AK15 residues, however, is somewhat different than that reported. In the top AF3 model, C85 is 12.8 Å and 7.1 Å, respectively, from H2AK13 and H2AK15, which varies somewhat from that of the reported cryo-EM structure (5.2 Å and 9.9 Å, respectively) (Figure 5C).

The histone H4 lysine 20 (H4K20) methyltransferase, SUV420H1 (KMT5B), is classified as both a PTM writer and a histone variant reader, since it binds to nucleosomes containing H2AZ. Binding of the H4 N-terminal tail to the SUV420H1 catalytic site (36) is correctly predicted in the AF3 model (Figure 5D), which is notable given that the training data set contained no examples of SUV420H1 bound to the H4 tail. We note that the H4 tail in the cryo-EM structure of SUV420H1 bound to a nucleosome containing H2AZ (36) contains a K20M substitution, which drives tight binding to the active site. In addition, the side chain interactions between SUV420H1 and the H2AZ/H2B acidic patch are correctly predicted to the single-residue level (Figure 5D), as are interactions with the DNA (SI Figure 2B).

### Histone methyl-lysine specificity can be predicted by AF3

Many of the highest confidence models were readers of histone methyl-lysine residues. In particular, AF3 was able to recapitulate the conserved structural recognition of methyl-lysine residues by the aromatic cage of methyl-lysine reader proteins (Figure 6A-B, D-E). Many methyl reader domains recognize methyl-lysine within a specific sequence context, which means that they must also contact residues flanking the methylated lysine residue (37–39). We asked whether AF3 could be used to predict the ability of a methyl-lysine reader to selectively bind the methylated lysine within the correct sequence context. To test this, we designed an *in silico* nucleosome interaction screen of methyl-lysine reader proteins whose binding to a panel of modified nucleosomes had been previously assayed (40, 41). The candidates selected were the PWWP domain of DNMT3A, which binds selectively to H3K36me2/3, and the PHD domain of BPTF, which binds to H3K4me3.

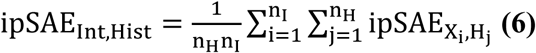

**Figure 6.**
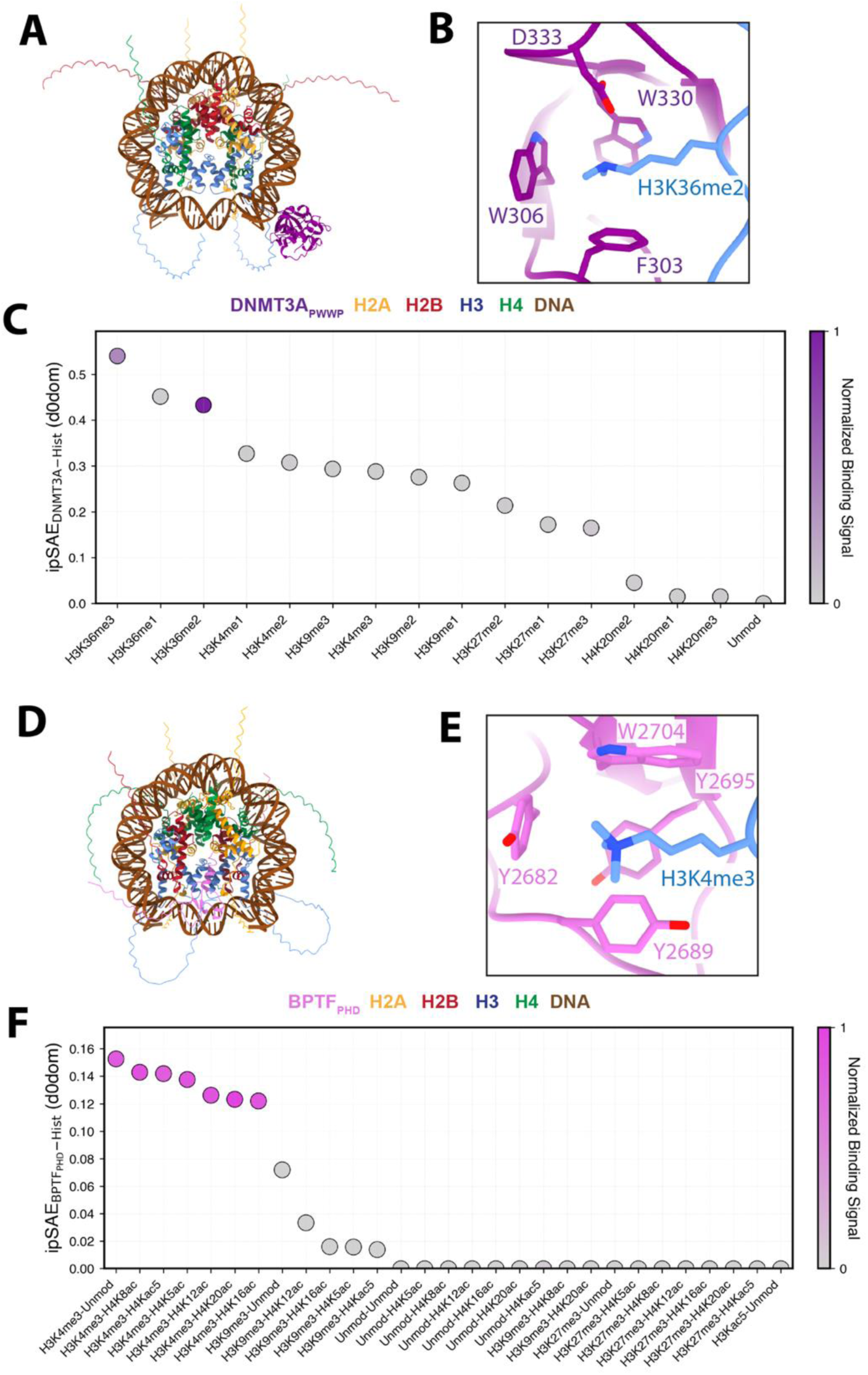
AlphaFold3 (AF3) can predict histone methyl-lysine specificity. (A) AF3 structure of DNMT3A_PWWP_ bound to a H3K36me2 nucleosome. (B) Zoom-in on the recognition of H3K36me2 by the DNMT3A_PWWP_ aromatic cage. (C) Distribution of the ipSAE score between DNMT3A_PWWP_ and the histone octamer (see Eq. 6) of modified nucleosomes tested in Ref. (40) predicted in AF3. (D) AF3 structure of BPTF_PHD_ bound to a H3K4me3 nucleosome. (E) Zoom-in on the recognition of H3K4me3 by the BPTF_PHD_ aromatic cage. (F) Distribution of the ipSAE score between BPTF_PHD_ and the histone octamer of modified nucleosomes tested in Ref. (41).

We performed AF3 predictions of each domain on the set of singly-modified (heterotypic) nucleosomes that had been assayed experimentally (40, 41) and scored models based on the confidence scores. Since the presence of disordered regions and/or accessory domains that do not contribute to protein binding negatively impacts the ipTM score (17, 25, 26) (SI Figure 1), we instead used the ipSAE (interface prediction Score from Aligned Errors), a metric that was developed to assess the confidence scores of biomolecule interactions in AF3 (26). ipSAE uses only the Predicted Aligned Error (PAE) values of residue pairs that are within a user-defined distance and PAE cutoff. Importantly, this approach has been shown to better separate true and false complexes over current existing metrics (26, 42). We scored each prediction using ipSAE, with distance and PAE cutoffs of 10 Å based on prior benchmarking datasets (26, 42). In addition, for ipSAE to reflect the confidence score between the interactor and the modified nucleosome, we calculated an average ipSAE score of the pairwise interactions between the interactor and the histone octamer which we define as ipSAE_Int,Hist_:

We excluded DNA chains from our averages, since all screened modifications are in the histone tails, so we would only expect to see changes in ipSAE scores between the interactor and the histone octamer. We calculated and extracted the highest ipSAE_Int,Hist_ across all five models for each prediction.

We first screened DNMT3A_PWWP_ against the panel of methylated nucleosomes and compared the ipSAE_Int,Hist_ scores with the reported normalized experimental binding signal (40).

DNMT3A_PWWP_ is an H3K36me2/3 reader that directs the recruitment of DNMT3A (40, 43, 44). In vitro, DNMT3A_PWWP_ has a higher affinity for H3K36me2 than for H3K36me3 (40). Across all predictions, the predicted structures of DNMT3A_PWWP_ and of the nucleosome match the crystal structures of the isolated entities (6, 45) (SI Figure 3). The complexes with mono-, di- and tri-methylated H3K36me nucleosomes all showed the methylated lysine inserted into the DNMT3A_PWWP_ aromatic cage and had the highest overall ipSAE_Int,Hist_ scores compared to the other predictions with modified nucleosomes (Figure 6C). We note, however, that the relative ranking by ipSAE_Int,Hist_ does not fully recapitulate the measured binding affinities. Whereas DNMT3A_PWWP_ has been shown to bind most tightly to H3K36me2, followed by H3K36me3, and exhibited no detectable binding to H3K36me1 nucleosomes (40), H3K36me3 has the highest ipSAE_Int,Hist_ score and scores somewhat higher than H3K36me2 (figure 6C).

AF3 predictions for the PHD domain of BPTF (BPTF_PHD_), which specifically recognizes H3K4me3 (46), were done with the same set of nucleosomes containing different combinations of methylated and acetylated histones that had been tested experimentally (41). BPTF is a ∼300kDa subunit of the Nucleosome Remodeling Factor (NURF) complex, an ATP-dependent chromatin remodeling complex that repositions nucleosomes in euchromatin (46). In a screen testing BPTF_PHD_ against a panel of modified nucleosomes, BPTF_PHD_ bound all nucleosomes containing H3K4me3 with similar affinity, irrespective of the presence or absence of other modifications (41). Our AF3 predictions yielded models of BPTF that match the previously resolved crystal structure and intact modified nucleosome structures (SI Figure 4). Based on ipSAE_Int,Hist_, AF3 was able to successfully rank all H3K4me3-containing nucleosomes with the highest confidence (Figure 6D-F), although we note that the range of confidence scores across all models were lower than those obtained in our virtual screen of DNMT3A_PWWP_. AF3 also correctly oriented H3K4me3 within the aromatic cage of BPTF_PHD_, matching the crystal structure of BPTF bound to a H3K4me3 peptide (47) (SI Figure 4C).

## Discussion

The development of AlphaFold3 represents a major advance in biomolecular structure prediction because of its ability to model interactions between proteins, nucleic acids, ions and small molecules, and its incorporation of post-translational modifications. This expanded capability makes AF3 a powerful tool for integration with chromatin structural biology. Our systematic and quantitative assessment of AF3’s performance in predicting nucleosome-containing complexes identified contexts in which AF3 performs remarkably well and scenarios in which its predictions are unreliable.

Our benchmarking reveals a clear dichotomy in AF3 performance between different modes of nucleosome recognition. AF3 excels at predicting complexes in which binding is driven primarily by interactions with the histone octamer, including histone variants, PTM writers, erasers, and readers (Figure 3C). In these cases, AF3 frequently recapitulates experimentally determined binding orientations with high accuracy and, in several instances, reproduces residue-level interactions that underlie PTM recognition and enzyme specificity (Figures 5 and 6).

Notably, AF3 correctly positions modified residues within reader domains and captures auxiliary interactions with the nucleosome acidic patch and DNA backbone, even when these interactions involve flexible histone tails or were not explicitly present in the training dataset. These results underscore AF3’s capacity to generalize learned biochemical principles of protein-protein interactions, including those involving modified side chains. By contrast, AF3 performs substantially worse for complexes in which nucleosome binding is dominated by protein-DNA interactions (Figure 4), particularly for transcription factors and chromatin remodelers. In these cases, predicted models frequently exhibit incorrect positioning of interactors along the superhelical locations of nucleosomal DNA, distorted DNA geometry, or physically unrealistic DNA topology.

Our findings highlight an important conceptual limitation of current deep learning-based structure prediction approaches. Whereas AF3 accurately captures protein-protein interfaces and histone-driven nucleosome recognition, it does not yet fully model the energetic and topological constraints governing protein engagement with nucleosomal DNA. For transcription factors and nucleosome remodelers, productive binding often involves numerous thermodynamic considerations such as competition with histone-DNA contacts and local deformation of DNA (48, 49), features that may be underrepresented or insufficiently parameterized in the training data. It is important to note that inaccuracies in predicted protein-DNA interactions are not confined to models of proteins bound to nucleosomes but reflect a broader challenge in structure prediction. A recent comprehensive biomolecular structure benchmark found protein-nucleic acid prediction to be one of the worst-performing tasks across all biomolecular structure prediction algorithms (50). Addressing these challenges will likely require advances in how nucleic acids are represented within structure prediction frameworks and a larger training dataset, as only <10% of current PDB entries contain nucleic acids.

This work introduces interactor-specific evaluation metrics that disentangle prediction accuracy of the nucleosome core from that of the nucleosome-binding factor. By partitioning DockQ into interactor-histone and interactor-DNA components, we show that conventional global metrics can substantially overestimate the accuracy of nucleosome-bound predictions due to the highly conserved and accurately predicted histone octamer. Our interactor-focused metrics provide a more faithful assessment of functional interfaces and should be broadly applicable to benchmarking other chromatin-associated complexes.

Beyond benchmarking against known structures, we demonstrate that AF3 can be leveraged to predict histone PTM specificity in silico. Using ipSAE-based scoring, AF3 successfully prioritizes cognate methyl-lysine marks for well-characterized reader domains, correctly ranking interactions with H3K36- and H3K4-methylated nucleosomes above other types of modifications (Figure 6). Although AF3 does not reliably distinguish between different methylation states of the same lysine, its ability to recover qualitative specificity suggests that AF3-based screening can complement biochemical assays by narrowing candidate PTM-reader interactions and informing experimental design. Evaluation of the ability of AF3 to predict reader interactions with PTMs other than methyl-lysine awaits experimental data on nucleosomes containing other types of modifications.

A major limitation of AF3 and, more generally, of biomolecular structure prediction algorithms is their inability to capture and consider contributions of molecular dynamics (51). Many nucleosome-binding proteins, especially those that recognize PTMs on histone tails, have binding affinities that are influenced by changes in nucleosome conformation and flexibility.

Histone modifications such as acetylation and ubiquitination can indirectly regulate nucleosome-binding interactions by altering the accessibility of histone tails or shielding critical nucleosome-binding interfaces (2, 52). We illustrate this limitation using a nucleosome interaction screen with the BPTF_PHD-BD_ module, which interacts bivalently with H3K4me3 and acetylated histone H4 (SI Figure 5). Previous work had found that, when the N-terminal of histone H4 is acetylated at multiple sites (H4Kac5), BPTF_PHD-BD_ pulls down H3K4me3-H4Kac5 with a 7-fold enrichment over other nucleosomes (41). This enhanced affinity is likely driven by increased histone tail accessibility and a higher frequency of binding events, rather than by direct accommodation of multiple acetylated lysines, since BPTF_BD_ can only bind one acetylated lysine (53). In our AF3-based screen, H3K4me3-H4Kac5 nucleosomes scored unexpectedly low in confidence compared to other modifications, even ranking below some H3K9me3-containing nucleosomes. This discrepancy underscores AF3’s inability to capture dynamic changes in nucleosome accessibility and conformational flexibility that influence binding.

Our benchmarking highlights both the strengths and weaknesses of AlphaFold3 in predicting structures of proteins bound to nucleosomes. AF3 consistently captures key features of nucleosome recognition, enabling the generation of mechanistic hypotheses that can guide experimental design. While challenges remain, particularly in modeling dynamic and DNA-driven interactions, AF3’s ability to reproduce critical nucleosome-interaction interfaces positions AF3 as a transformative tool for chromatin biology.

## Methods

### Benchmark dataset curation

To construct a benchmark set of nucleosome-containing complexes, we queried the Protein Data Bank (PDB) for entries containing a nucleosome or nucleosome-like particle deposited after the AlphaFold3 (AF3) training dataset cutoff date (September 30^th^, 2021) up until September 1, 2024 (Figure 1A) (19). To reduce potential training-set leakage through post-cutoff redepositions, we also removed post-cutoff entries for which the same structure had been released prior to the cutoff.

Beginning with the post-cutoff pool (n=322), we applied a series of quality and compatibility filters. We removed complexes whose total number of amino acids, DNA bases, ligands, modifications, and ions exceeded the AF3 server input limit of 5,000 tokens. Complexes containing explicitly modeled DNA damage or chemically modified nucleotides were also removed. To remove redundancy from the evaluation set, we collapsed PDB entries into unique complexes by comparing protein sequences, nucleic acid sequences, and stoichiometry. PDB entries were considered redundant if they contained the same sequences for all protein and nucleic acid chains and chain stoichiometry. When multiple redundant entries existed, we retained a single representative structure, selecting the complex that was deposited first. After filtering, this yielded 115 unique nucleosome-containing complexes used throughout the benchmark that were successfully run on AF3 or AF3X (Figure 1B).

### Incorporation of post-translational modifications, ligands, and ions

Supported post-translational modifications (e.g., common lysine methylation and acetylation marks on histone tails) were specified explicitly in the AF3 JSON input. Structures containing a covalently linked ubiquitin protein were predicted using AlphaFold3X (AF3X), which allows specification of covalent bonds (22). Ligands and ions not supported by the AF3 release were removed or replaced when possible, with the closest chemically compatible AF3 representation (SI Table 1).

### AlphaFold3 and AlphaFold3x prediction parameters

All structure predictions were generated using the publicly available AF3 server with default inference parameters, with the exception of structures containing ubiquitin. Predictions were performed using sequences assigned to the PDB entry. Under default settings, the server performs ten recycling iterations during structure refinement and generates five independent models, all of which were retained for downstream analysis. All predictions were performed with random seed numbers and template searching was enabled using the default PDB cutoff date (September 30^th^, 2021).

Because the AF3 server does not support ubiquitin covalently linked to lysine via an isopeptide bond, ubiquitinated nucleosomes were modeled using AlphaFold3X (AF3X) (22). For each ubiquitin-containing complex, we encoded a crosslink between the C-terminal residue of ubiquitin (G76) and the sidechain of the modified histone lysine (e.g., H2AK13 or H2AK15) by substituting each residue with alanine and encoding a disulfide bond between the two residues. All AF3X structure predictions were performed on a local workstation. Each prediction was performed with a random seed number and ten recycles. Multiple sequence alignments were generated using HMMER v3.4-based searches against the AF3 public sequence databases.

### DockQ scoring, iRMSD, and Interactor-Nucleosome calculations

Interface accuracy was quantified using DockQ (28, 29), which combines the fraction of native contacts (f_nat_), interface RMSD (iRMSD), and ligand RMSD (LRMSD) into a single score ranging from 0 to 1. Native contacts were defined using heavy-atom pairs within 5 Å in the reference structure and iRMSD was computed over interface residues following optimal superposition. For multi-chain assemblies, global DockQ was calculated as the unweighted mean across all interacting chain pairs satisfying the DockQ contact criterion and is reported as the “overall DockQ” metric (Figure 2).

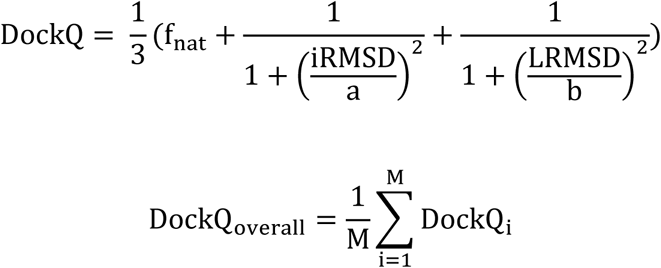

For each prediction, we computed DockQ scores between each of the five AF3 (or AF3X) models and the corresponding experimental structure. For complexes containing at least one interactor, chains were assigned to three groups: histones (all H2A/H2B/H3/H4 chains and variants), DNA, and interactors (all non-histone, non-DNA chains).

Pairwise DockQ values between interactor-histone (DockQ_Int,Hist_), interactor-DNA (DockQ_Int,DNA_), and interactor-nucleosome (DockQ_Int–Nuc_) chain pairs were defined as:

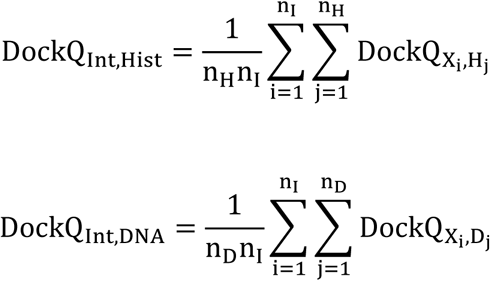

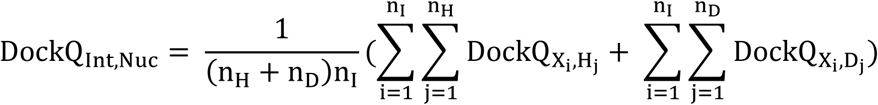

For multi-nucleosome complexes, severe clashes between DNA and histone cores prevented reliable alignment, so these models were excluded from any DockQ analysis.

### Post Translational Modification (PTM) screen

All predictions of proteins bound to histones containing post-translational modifications (PTMs) were carried out on the AF3 public webserver using default settings, generating five models per prediction. We used the same isolated reader fragments as the experimental studies (DNMT3A_PWWP_ and BPTF_PHD_) and retained the same fragment boundaries as reported (40, 41). These domains were screened against panels of heterotypic nucleosomes matching the modification sets used in the corresponding experimental screens.

Predictions were ranked using ipSAE (interface prediction Score from Aligned Errors), which reweights AF3 confidence by restricting scoring to residue pairs that satisfy user-defined geometric and error constraints. The ipSAE metric has been shown to improve separation of true versus false interactions in prior benchmark studies (26, 42). Specifically, we used the ipSAE d0dom formulation as defined in the ipSAE method (26) and applied a distance cutoff of 10 Å and a PAE cutoff of 10 Å.

Since PTM recognition of DNMT3A_PWWP_ and BPTF_PHD_ is driven by histone-mediated interactions we scored models based on ipSAE_Int,Hist_. This is computed by averaging pairwise ipSAE values between the interactor and histone chains. This is defined as:

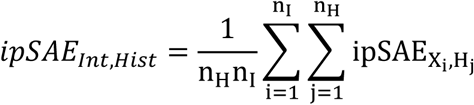

For each prediction, we took the maximum ipSAE_Int,Hist_ across all five AF3 predictions.

## ACKNOWLEDGEMENTS

We thank Dr. Jeffrey Gray for advice and critical feedback. Supported by the National Institutes of Health under National Institute of General Medical Sciences award R35GM130393 (C.W.) and National Cancer Institute fellowship F31CA271243 (S.R.).

## DATA AVALIABILITY

All AlphaFold3 and AlphaFold3X predictions, DockQ outputs, ipSAE output, and analysis scripts are available on Zenodo (10.5281/zenodo.20737868).

## SUPPLEMENTAL INFORMATION

**SI Figure 1.**
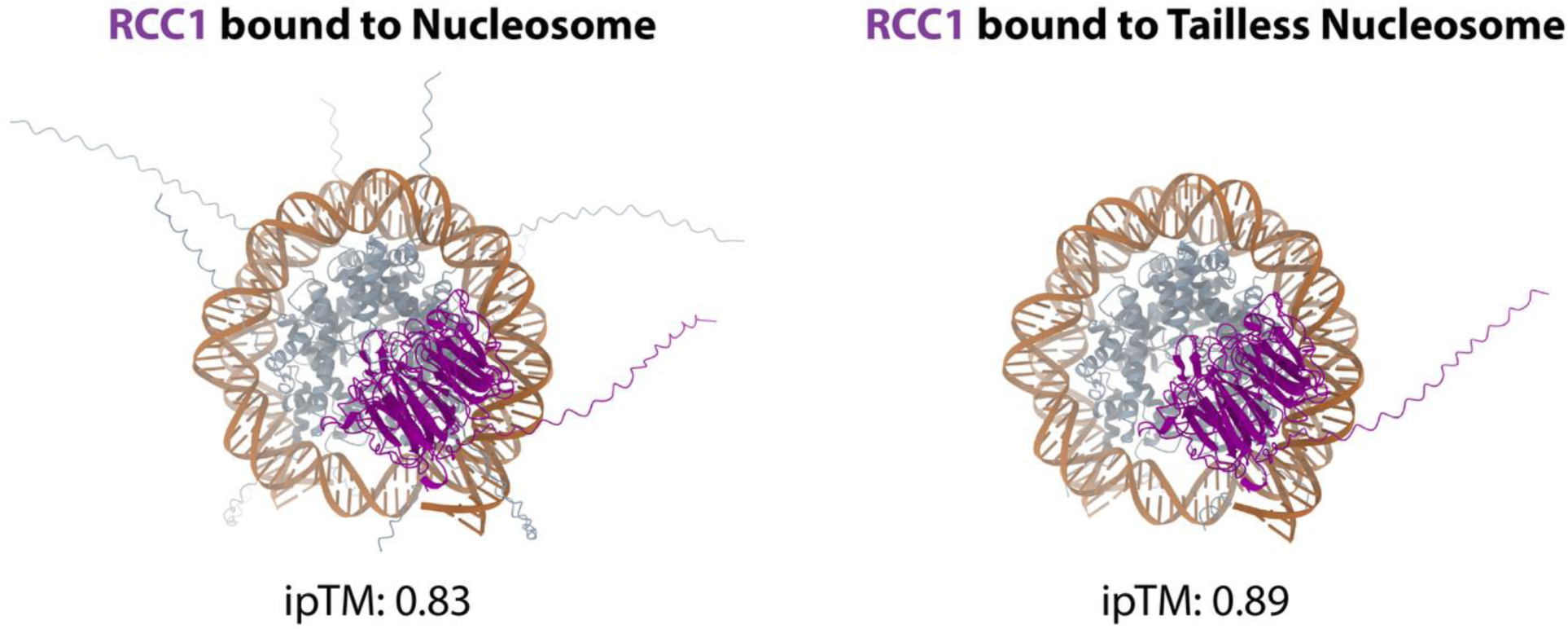
AlphaFold3 model of RCC1 with unmodified and tailless nucleosomes. The calculation of the overall interface predicted Template Modeling (ipTM) incorporates confidence values for all residues within a model. Therefore, the presence of non-interacting disordered regions, negatively impacts the ipTM score.

**SI Figure 2.**
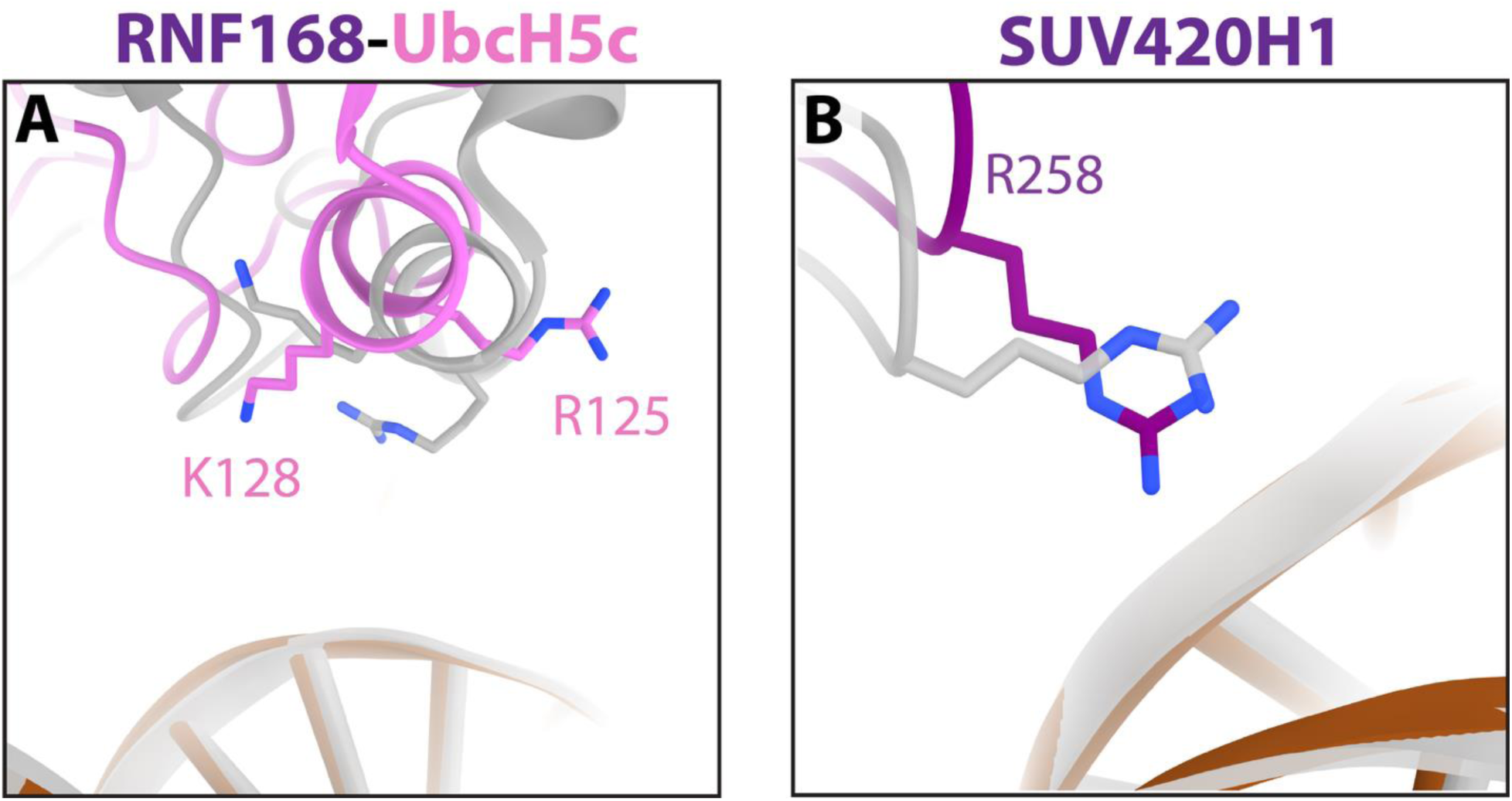
UbcH5c and SUV420H1 interactions with the nucleosome DNA. AlphaFold3 (AF3) models of (A) RNF168-UbcH5c and (B) SUV420H1 with a close-up view on the nucleosome DNA interactions. The AF3 model (colored) is superimposed onto the experimental structure (gray).

**SI Figure 3.**
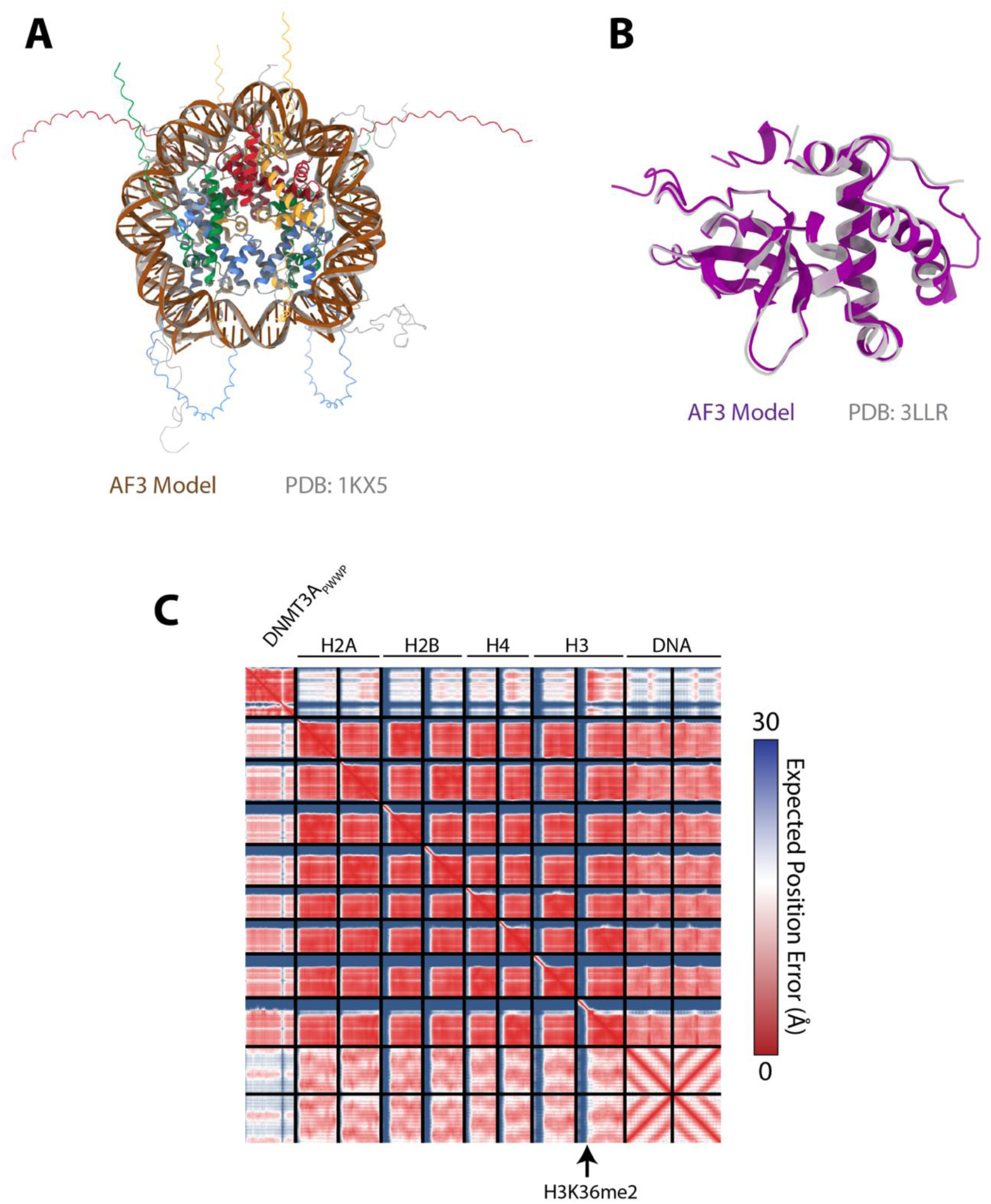
Structure prediction accuracy of DNMT3A_PWWP_ bound to H3K36me2 nucleosome. Comparison of AF3-predicted structures of the (A) H3K36me2 nucleosome and (B) DNMT3A_PWWP_ domain from the DNMT3A_PWWP_-H3K36me2 nucleosome model, superimposed onto the crystal structures of an unmodified nucleosome and DNMT3A_PWWP_, respectively. (C) Predicted Aligned Error of the top ipSAE-scoring model of DNMT3A_PWWP_ bound to a H3K36me2 nucleosome.

**SI Figure 4.**
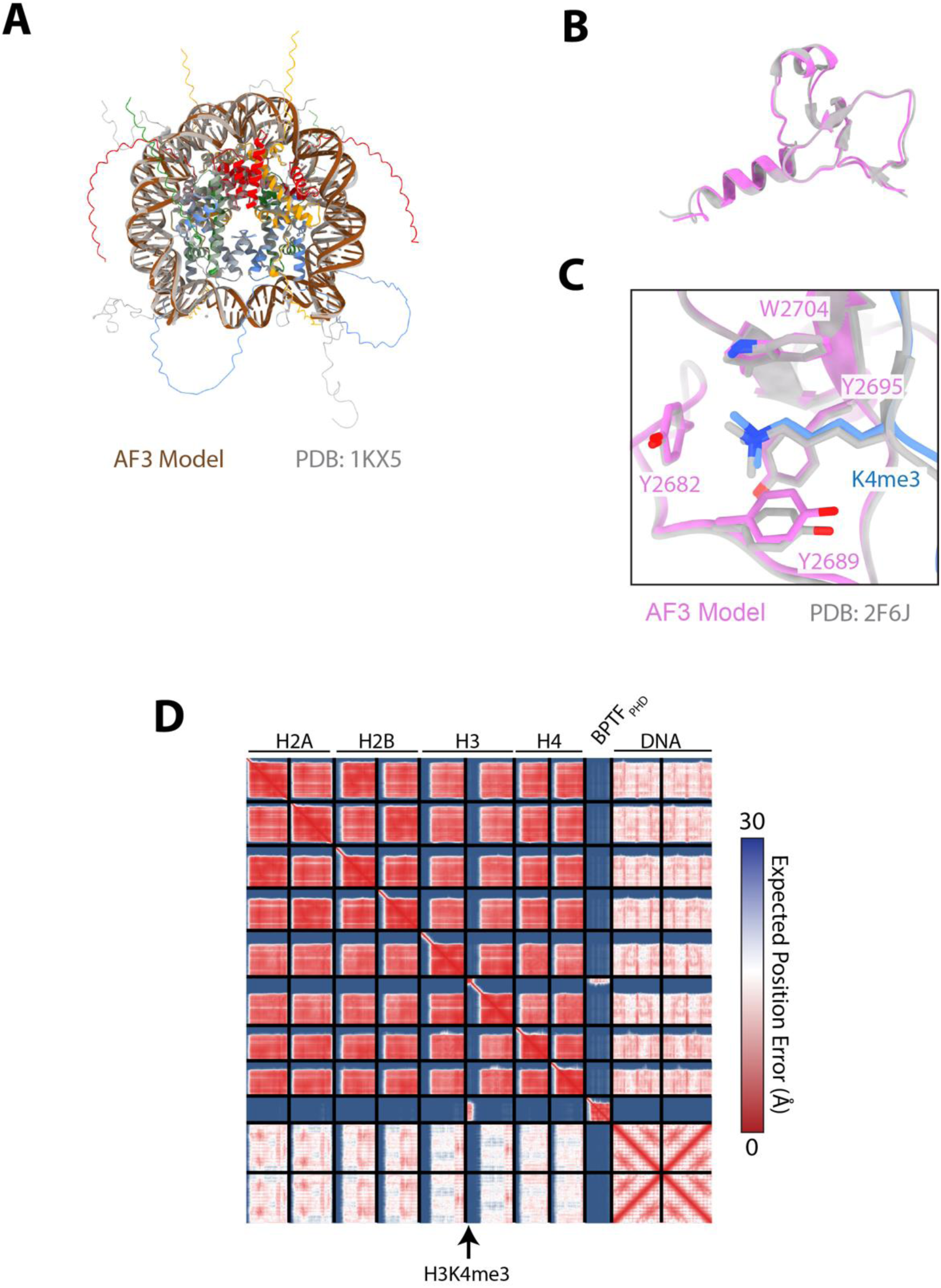
Structure prediction accuracy of BPTF_PHD_ bound to H3K4me3 nucleosome. Comparison of AF3-predicted structures of the (A) H3K4me3 nucleosome and (B) BPTF_PHD_ domain from the BPTF_PHD_- H3K4me3 nucleosome model, superimposed onto the crystal structures of an unmodified nucleosome and BPTF_PHD_, respectively. (C) Comparison of the interaction between H3K4me3 and BPTF_PHD_ in the AF3 model and crystal structure (47). (D) Predicted Aligned Error of the top ipSAE-scoring model of BPTF_PHD_ bound to a H3K4me3 nucleosome.

**SI Figure 5.**
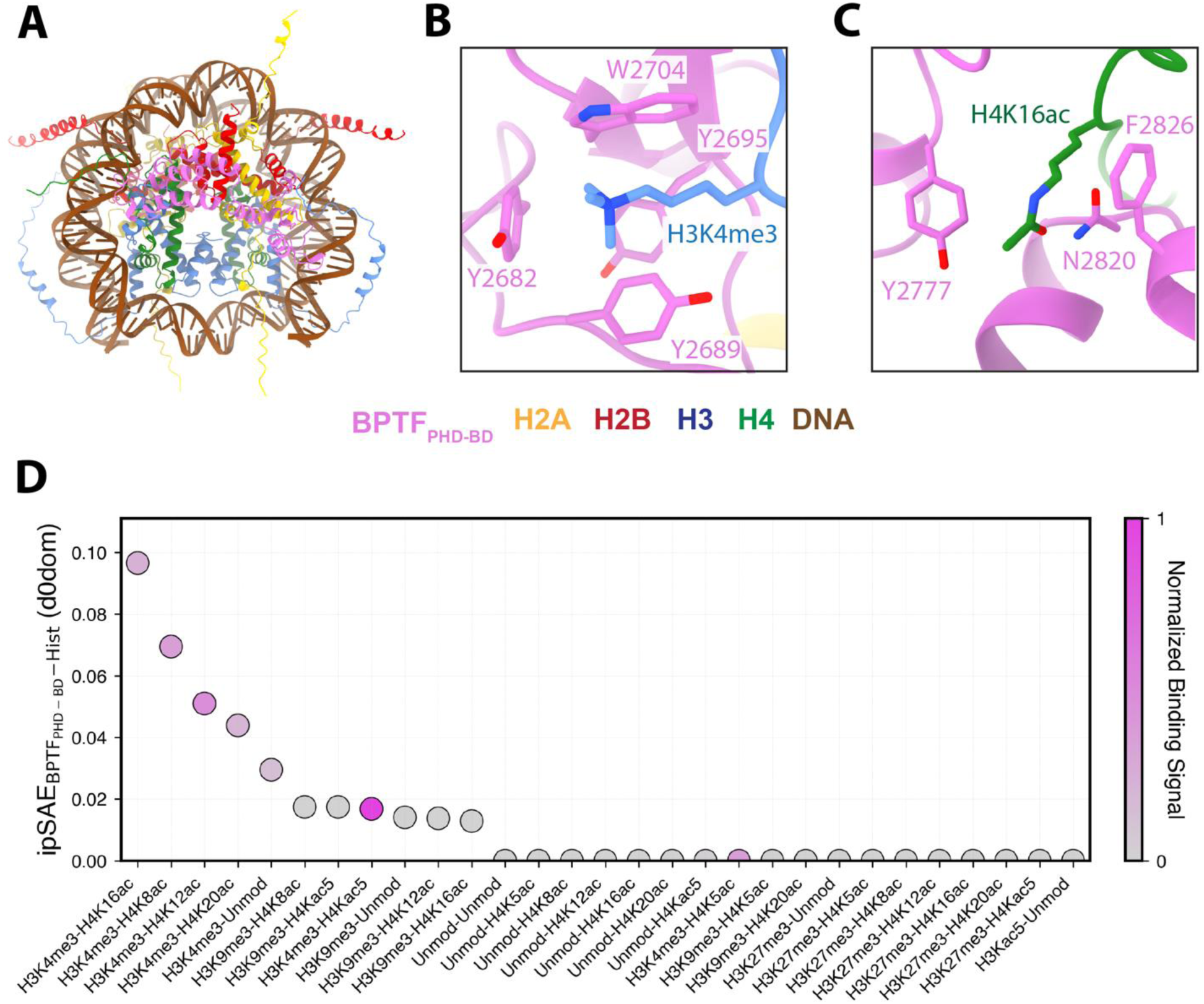
(A) AF3 structure of BPTF_PHD-BD_ bound to a H3K4me3 nucleosome. (B) Zoom-in on the recognition of H3K4me3 by the BPTF_PHD_ aromatic cage. (C) Zoom-in on the recognition of H4K16ac by the BPTF_BD_ domain. (D) Distribution of the ipSAE score between BPTF_PHD-BD_ and the histone octamer of modified nucleosomes tested in Ref. (41).

**SI Table 1.**
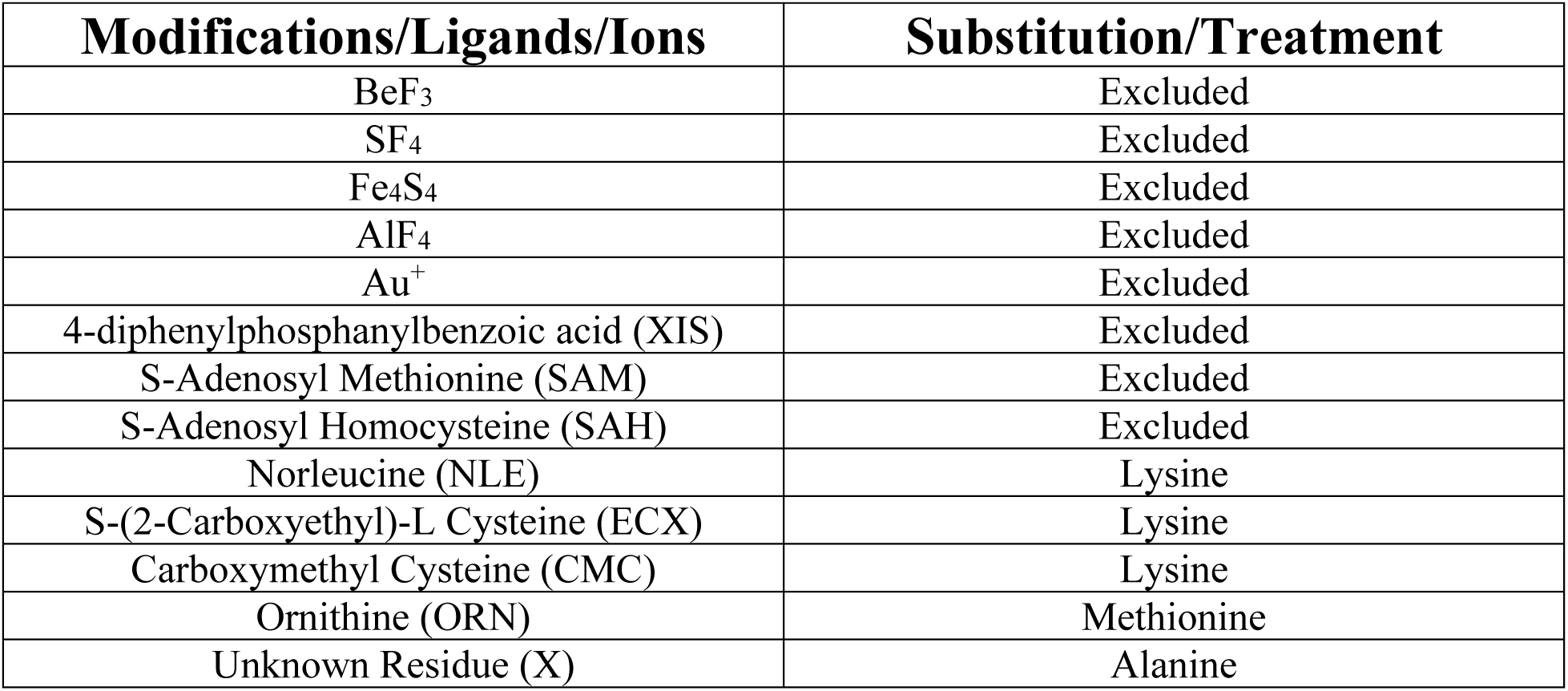
Treatment and substitutions of modifications, ligands, and ions not supported in AF3.

