## Supplemental Figures for "A Chromatin Biology Assessment of AlphaFold3"

**SUPPLEMENTAL INFORMATION**

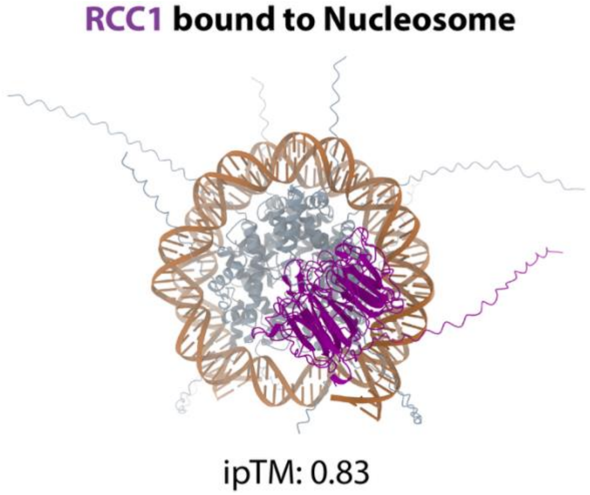


**SI Figure 1. AlphaFold3 model of RCC1 with unmodified and tailless nucleosomes.** The calculation of the overall interface predicted Template Modeling (ipTM) incorporates confidence values for all residues within a model. Therefore, the presence of non-interacting disordered regions, negatively impacts the ipTM score.

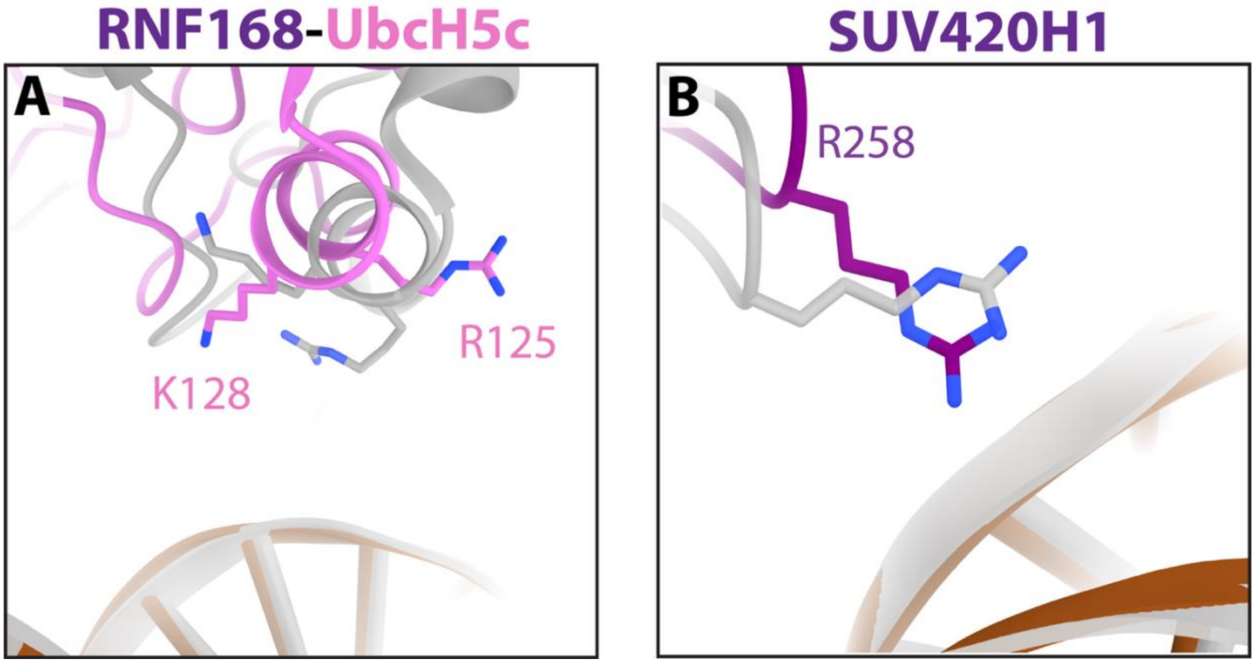


**SI Figure 2. UbcH5c and SUV420H1 interactions with the nucleosome DNA.** AlphaFold3 (AF3) models of (A) RNF168-UbcH5c and (B) SUV420H1 with a close-up view on the nucleosome DNA interactions. The AF3 model (colored) is superimposed onto the experimental structure (gray).

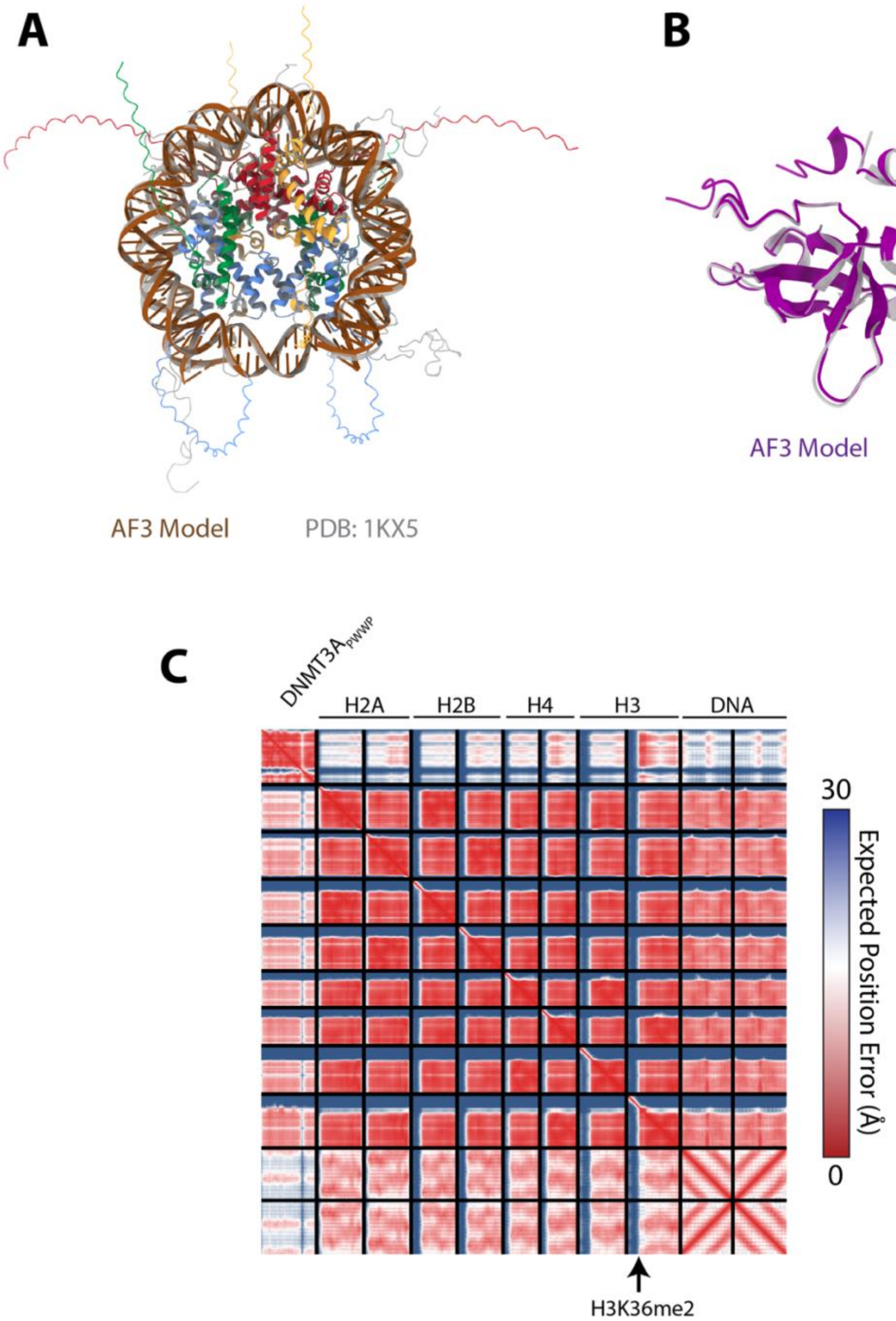


**SI Figure 3. Structure prediction accuracy of DNMT3A$_{PWWP}$ bound to H3K36me2 nucleosome.** Comparison of AF3-predicted structures of the (A) H3K36me2 nucleosome and (B) DNMT3A$_{PWWP}$ domain from the DNMT3A$_{PWWP}$-H3K36me2 nucleosome model, superimposed onto the crystal structures of an unmodified nucleosome and DNMT3A$_{PWWP}$, respectively. (C) Predicted Aligned Error of the top ipSAE-scoring model of DNMT3A$_{PWWP}$ bound to a H3K36me2 nucleosome.

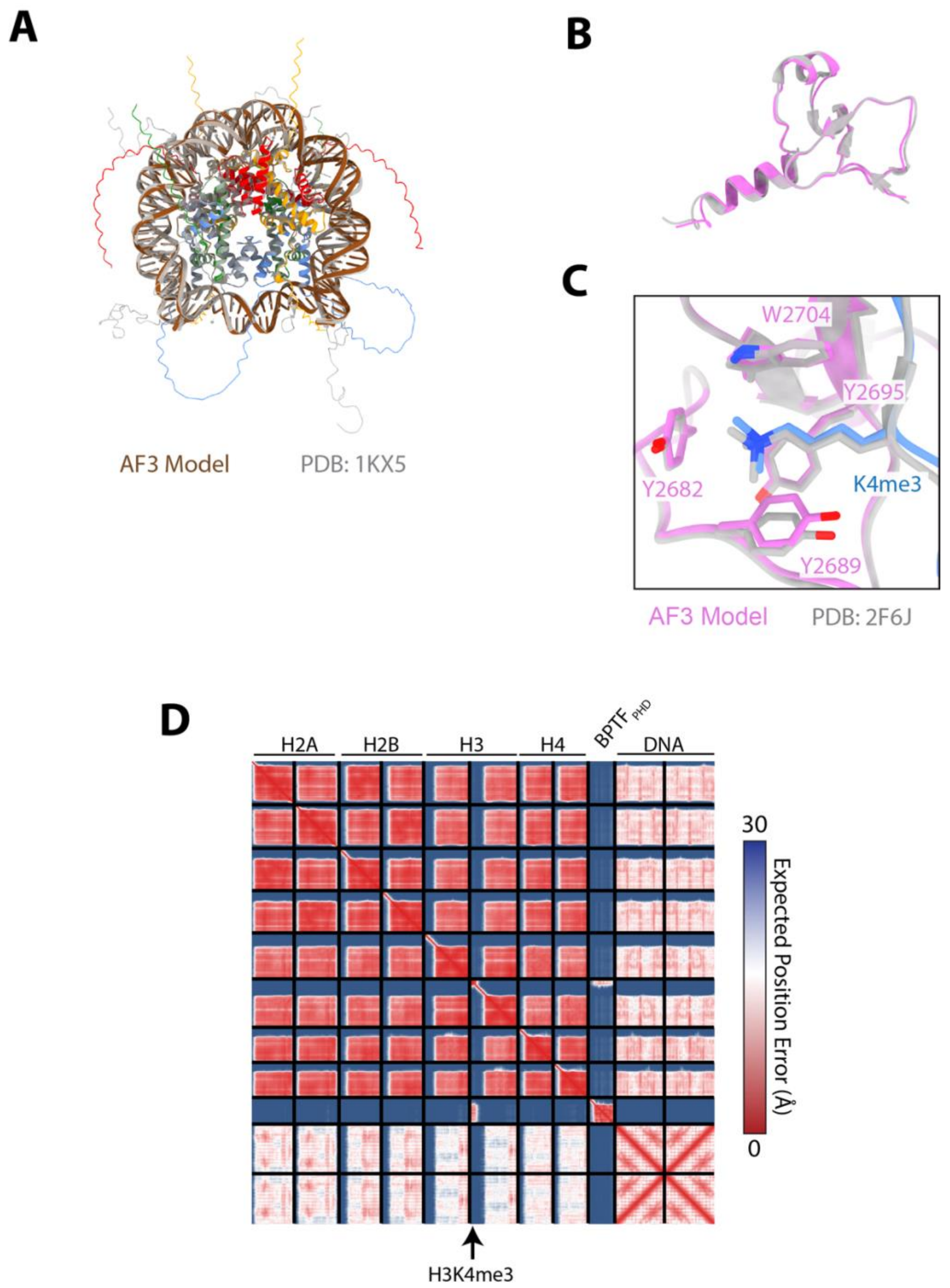


**SI Figure 4. Structure prediction accuracy of BPTF$_{PHD}$ bound to H3K4me3 nucleosome.** Comparison of AF3-predicted structures of the (A) H3K4me3 nucleosome and (B) BPTF$_{PHD}$ domain from the BPTF$_{PHD}$- H3K4me3 nucleosome model, superimposed onto the crystal structures of an unmodified nucleosome and BPTF$_{PHD}$, respectively. (C) Comparison of the interaction between H3K4me3 and BPTF$_{PHD}$ in the AF3 model and crystal structure (47). (D) Predicted Aligned Error of the top ipSAE-scoring model of BPTF$_{PHD}$ bound to a H3K4me3 nucleosome.

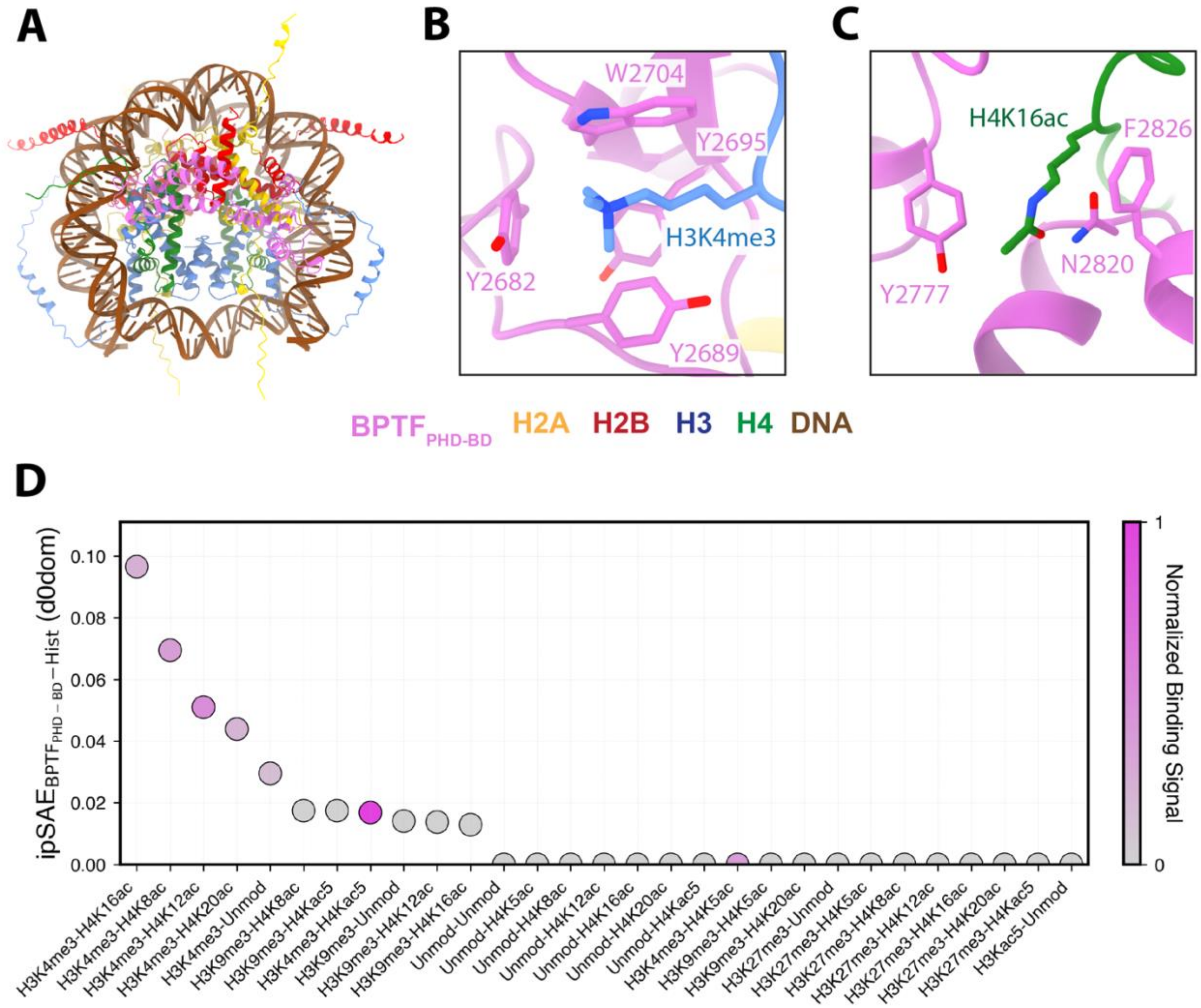


**SI Figure 5.** (A) AF3 structure of $BPTF_{PHD-BD}$ bound to a H3K4me3 nucleosome. (B) Zoom-in on the recognition of H3K4me3 by the $BPTF_{PHD}$ aromatic cage. (C) Zoom-in on the recognition of H4K16ac by the $BPTF_{BD}$ domain. (D) Distribution of the ipSAE score between $BPTF_{PHD-BD}$ and the histone octamer of modified nucleosomes tested in Ref. (41).

**SI Table 1. Treatment and substitutions of modifications, ligands, and ions not supported in AF3.**

| Modifications/Ligands/Ions | Substitution/Treatment |
|---|---|
| $BeF_3$ | Excluded |
| $SF_4$ | Excluded |
| $Fe_4S_4$ | Excluded |
| $AlF_4$ | Excluded |
| $Au^+$ | Excluded |
| 4-diphenylphosphanylbenzoic acid (XIS) | Excluded |
| S-Adenosyl Methionine (SAM) | Excluded |
| S-Adenosyl Homocysteine (SAH) | Excluded |
| Norleucine (NLE) | Lysine |
| S-(2-Carboxyethyl)-L Cysteine (ECX) | Lysine |
| Carboxymethyl Cysteine (CMC) | Lysine |
| Ornithine (ORN) | Methionine |
| Unknown Residue (X) | Alanine |
